# Familial relationships in electronic health records (EHR) v2

**DOI:** 10.1101/731976

**Authors:** Zhouzerui Liu, Nicholas Tatonetti

## Abstract

Heritability is an important statistic for evaluating genetic contribution to phenotypes. Estimating heritability, however, requires a laborious recruitment of a large number of relatives. Electronic health records (EHR) contain massive relative information in emergency contact forms. Recently, we presented RIFTEHR, an algorithm for extracting relationships from EHR. Here, we present an updated version and reconstructed 4.2 million familial relationships from the latest New York-Presbyterian/Columbia University Irving Medical Center (CUIMC) EHR system. The number of updated relationships is 30 percent more than the last version. We present a new implementation of RIFTEHR, which runs in linear time, thus largely improves the speed of the algorithm. We also present a data encryption method, to protect patient privacy in running the algorithm. These resources can be used for generalized use of familial relationships from EHR in genetic studies.

## Introduction

Heritability is the fraction of phenotype variance due to genetic differences (Tenesa & Haley, 2013). It is valuable in identifying disease risk factors, predicting disease risk and preventing disease (Chatterjee, Shi, & García-Closas, 2016). The widely used method for estimating heritability without genetic data is family-based studies, especially twin studies. Family-based studies, however, require laborious recruitments of a large number of relatives, and are often limited by their sample sizes. The probable largest single study contains 203,691 twin individuals (Mucci et al., 2016), and the meta-analysis using 2,748 studies from 1955 includes 14.5 million people (Polderman et al., 2015).

Electronic health records (EHR) store massive patient relationships in their emergency contact forms. This information has not been used for genetic studies until recently. In previous work, we successfully reconstructed familial relationship networks of three medical centers using emergency contact information in EHR (Polubriaginof et al., 2018). This study involved in total 1.9 million subjects, which was much larger than any former single family-based study. This holds the promise that EHR can be utilized for large scale relative recruitment for genetic studies.

Relationship inference from the electronic health record (RIFTEHR) algorithm is used for extracting familial relationships from EHR (Polubriaginof et al., 2018). It maps emergency contact person to patients in the hospital database, and infers relationship networks and families. This algorithm can be used within a single medical center, or scaled up to integrate data from multiple institutions. But the current implementation of the mapping step runs in 𝒪(*n*^2^) time, thus may not be suitable for process extremely large-scale data. Also, the current algorithm can only be run by people with access to the identifiable data. This limits the usage of the algorithm.

In this study, we applied RIFTEHR algorithm to the latest CUIMC EHR system, and reconstructed 4.2 million familial relationships. To adapt the algorithm for large-scale data processing, we came up with a linear time implementation of mapping. And to protect patient privacy, we provided a data encryption method.

## Methods

### Datasets

The emergency contact and patient demographics dataset were collected from Columbia University Irving Medical Center EHR system. 823,112 patients were involved in the analysis.

### Software

Python 2.7

Julia 0.4.7

MySQL 5.7.26

### Extracting relationships from EHR

Enterprise Master Patient Index (EMPI), when available or the medical record number (MRN) is used as patient identifiers. Relationships were extracted using the RIFTEHR algorithm (Polubriaginof et al., 2018).

### Mapping emergency contact to patients

The original Python implementation was described before (Polubriaginof et al., 2018). The new implementation preprocesses data in Python, such as lower case converting for names, and finds exact matches for first name, last name, phone number, zip code, or their combinations using MySQL. The output of the new implementation is the same with the original one, including patient identifier, relationship, emergency contact identifier and the unique match combinations.

### Execution time of mapping

0.5, 1, 2 or 4 percent of 1.6 million emergency contact and 5.8 million patient demographics data from CUIMC EHR system were randomly selected as input datasets of the algorithm. Each dataset was fed to both implementations. The processor execution time of Python scripts was calculated using the time package. And the processor execution time of each SQL query was returned automatically in MySQL. Time for all scripts and queries were added up as the total execution time.

### Data encryption for patient privacy

The algorithm takes in the identifiable data, and output the according encrypted patient identifier, first name, last name, phone number and zip code data. A patient identifier and its encryption mapping table is also generated by the algorithm, for analysis which needs linkage to phenotypes in EHR. This encryption should be done before using RIFTEHR.

## Results

### Updated familial relationships

In the following context, we refer to results from (Polubriaginof et al., 2018) as the original or version 1, and refer to results from this study as the updated or version 2.

We retrieved emergency contact data from the latest CUIMC EHR and mapped it to the patient demographics database. In the updated database, 1,254,790 patients provided 1,643,011 emergency contact persons, and 5,873,115 patient demographics data are recorded. It is approximately 20 percent more than the original, which contains 1,353,045 emergency contact persons and 4,788,982 patient demographics.

Among 1.6 million emergency contact persons, we successfully mapped 1,297,768 to patients in the database as provided relationships. Of these, 589,152 are relatives. Utilizing provided relationships, we inferred additional 3,698,274 familial relationships. The total number of provided and inferred relationships is 4,287,426, which is 32 percent more than the original. (Table 1)

**Table 1.**
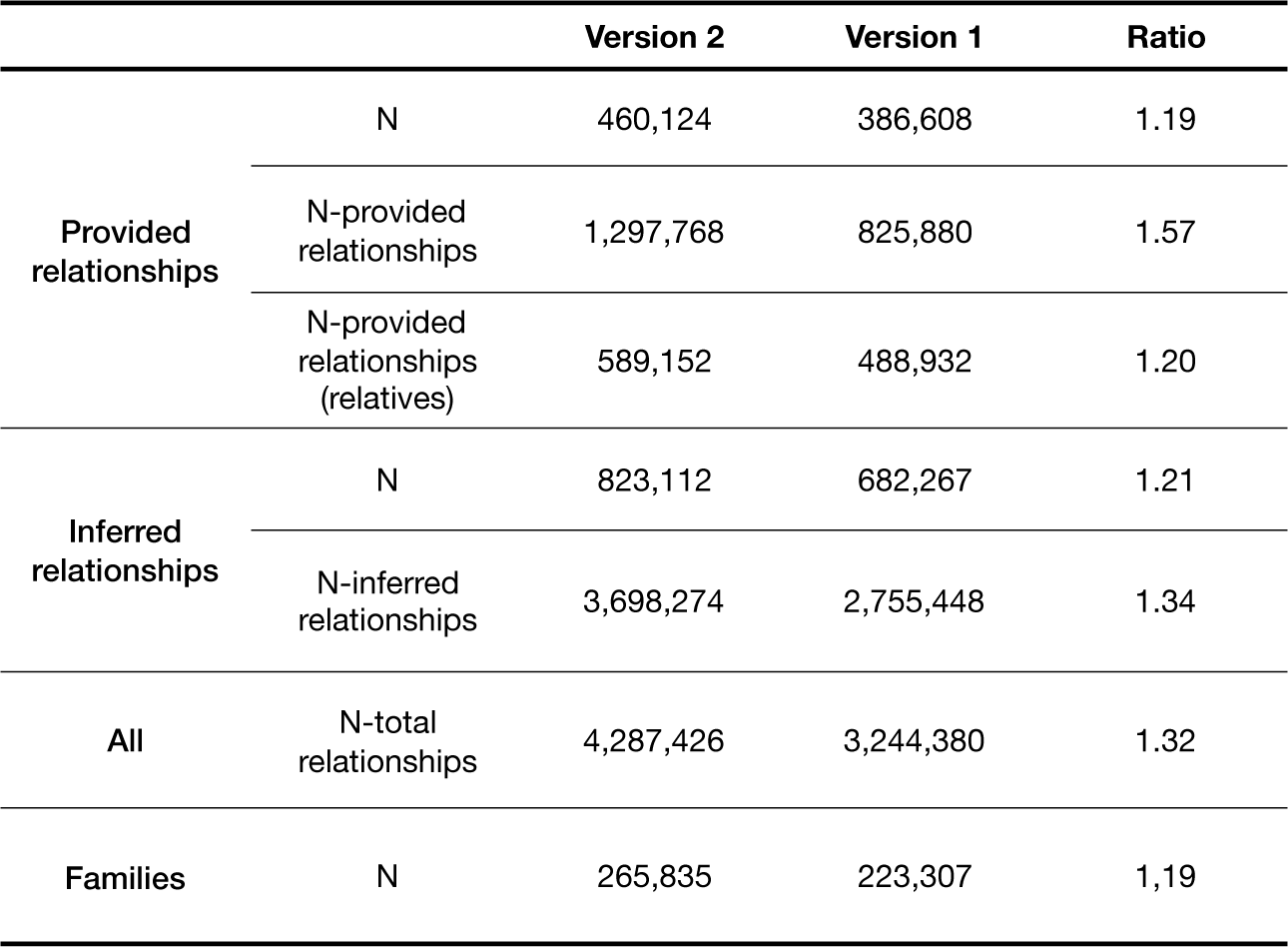
familial relationships extracted from EHR. Number of patients (N) involved in the relationships, Number of provided and inferred relationships and number of families are compared between the updated relationships (version 2) and the original version (version 1). Ratio is calculated as version 2/ version 1.

We then grouped patients into 265,835 families. (Table 1) The family size ranges from 2 to 231, which is much larger than the original. The maximum family size is 134 in version 1.

We further examined the degree of relationships. We got more first-degree relatives, but less second-, third- and fourth-degree relatives than version 1. (Table 2)

**Table 2.** degree of relationships. The number of first-, second-, third- and fourth-degree relative pairs is compared between 2 versions, as well as number of not blood relatives and unknown relatives. Ratio is calculated as version 2/ version 1.

|  |  | Version 2 | Version 1 | Ratio |
| --- | --- | --- | --- | --- |
|  | Twins | 14,588 | 15,616 | 0.93 |
| First<br>(e.g., child) | Siblings | 535,940 | 428,712 | 1.25 |
|  |  | 1,760,280 | 1,388,858 | 1.27 |
|  | Second<br>(e.g., grandchild, Nephew/Niece) | 797,346 | 825,880 | 0.97 |
|  | Third<br>(e.g., great-grandparent, cousin) | 319,524 | 488,932 | 0.65 |
|  | Fourth<br>(e.g., great-great-grandchild) | 552,358 | 682,267 | 0.81 |
|  | None<br>(e.g., spouse, in-laws) | 248,010 | 172,158 | 1.44 |
|  | Unknown<br>(e.g., parent/parent-in-law) | 595,215 | 429,880 | 1.38 |

### The new implementation of RIFTEHR

RIFTEHR algorithm extracted relationships by mapping emergency contact information to patient demographics data. The original Python implementation of the mapping uses brute-force search. Assuming the number of emergency contact entries is in direct ratio with the number of patients in the database, the complexity of the algorithm is 𝒪(*n*^2^). Considering the potential of data increasement in EHR, it will take a long time to execute. In this study, mapping 1.6 million emergency contact entries to 5.8 million patients takes approximately 2 days, if using 30 threads. To overcome the speed limitation of the algorithm, we therefore utilize SQL for mapping.

To compare the performance of two implementations, the execution times of various amount of data are calculated. Emergency contact and patient demographics data are random selected at a fixed percentage (0.5, 1, 2 or 4) of the total data, and the same dataset is mapped using both implementations. Compared to the original implementation, the new one significantly reduces the execution time. (Figure 1) The complexity of the new algorithm is 𝒪(*n*).

**Figure 1.**
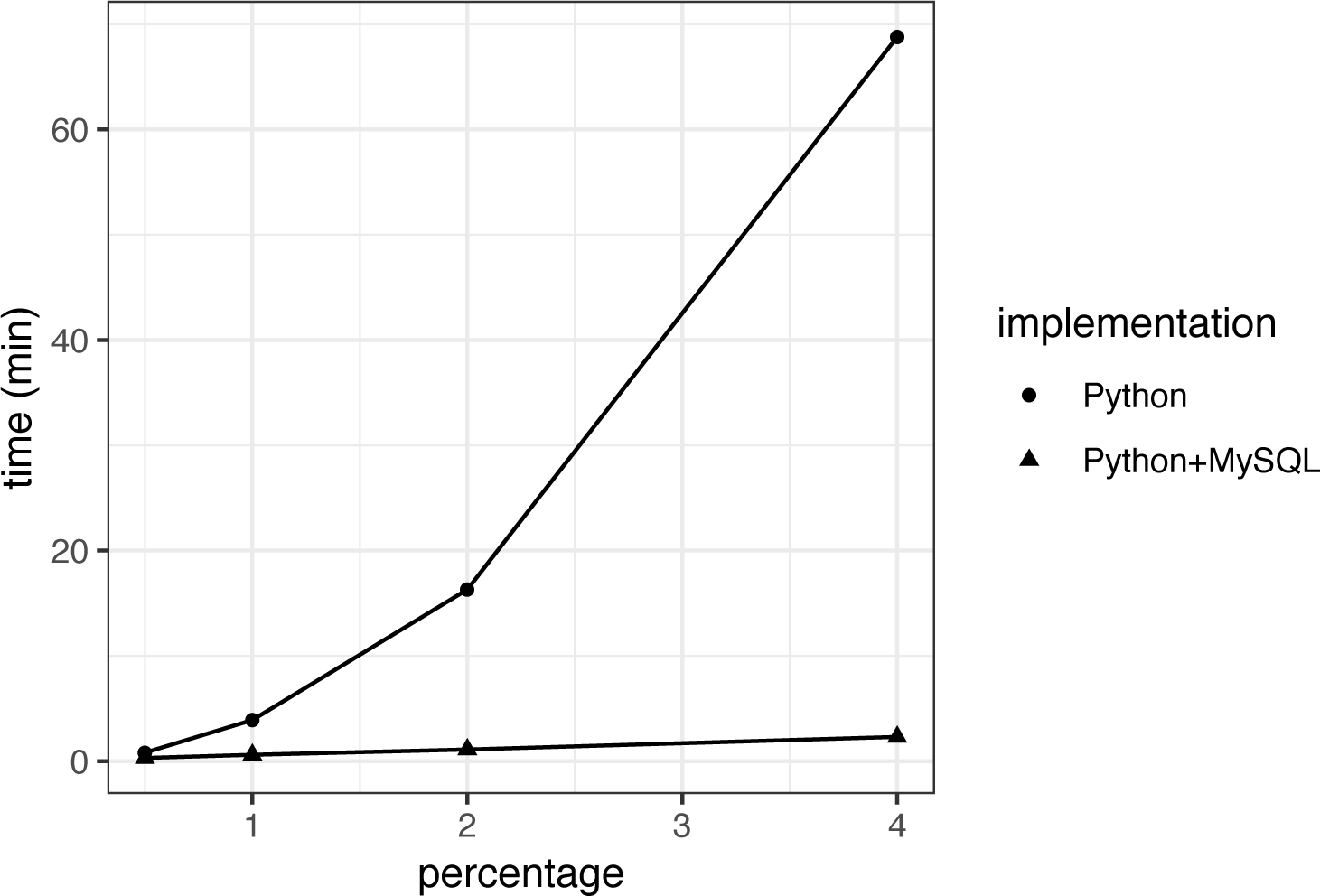
execution time of two implementations. The original implementation is in Python, and the new implementation uses Python and MySQL. 0.5, 1, 2 or 4 percent of 1.6 million emergency contact and 5.8 million patient demographics data is used as input of the algorithm. The execution time of the new implementation is significantly less than the original one, especially for large datasets.

### Data encryption for patient privacy

For those do not have access to identifiable data in EHR, we provided a data encryption method. The method is used before running RIFTEHR to encrypt patient identifier, first name, last name, phone number and zip code. Thus, no identifiable data is needed when extracting familial relationships from EHR using RIFTEHR. For studies need to link patient with data other than first name, last name, phone number and zip code in EHR, we also generated a patient identifier and its encryption mapping table during this data encryption.

## Discussion

Heritability is commonly estimated by family-based studies. Sample size is one of the key issues of these estimations. Here, we again presented the value of EHR as a comprehensive resource to overcome limitations in genetic studies. Leveraging the updated data from CUIMC EHR system and the RIFTEHR algorithm, we built over 4.2 million familial relationships and 260,000 family pedigrees from a single institution – a 30 percent increase than our previously study (Polubriaginof et al., 2018).

Besides twins and siblings, we also construct the extended family, including second-, third- and fourth-degree relatives. The distributions of relative degrees from the updated and original CUIMC EHR data show some differences. Version 2 has an increased number of first-degree, none and unknown relatives, and decrease in twins, second-, third- and fourth-degree relatives. Some decrease in number of twins is due to the integration of identifier numbers of the same patient. Decrease in second-, third- and fourth-degree relatives, and increase in none and unknown relatives, suggest we probably filtered out many non-blood relatives, and the data is cleaner than the original.

The EHR is a valuable source of phenotype information for scientific and health research. Family relationships already captured by the EHR can be extracted and used to estimate important disease parameters, like sibling co-occurrence and heritability. Here, we improve on the previously published extraction algorithm. We created a linear time implementation of RIFTEHR algorithm which will better scale with the data as it continues to grow. We also provided a demographic information encryption method, to protect patient privacy in extracting familial relationships.

## References

Chatterjee, N., Shi, J., & García-Closas, M. (2016). Developing and evaluating polygenic risk prediction models for stratified disease prevention. Nature Reviews Genetics, 17(7), 392.

Mucci, L. A., Hjelmborg, J. B., Harris, J. R., Czene, K., Havelick, D. J., Scheike, T., … Unger, R. H. (2016). Familial risk and heritability of cancer among twins in Nordic countries. Jama, 315(1), 68–76.

Polderman, T. J. C., Benyamin, B., De Leeuw, C. A., Sullivan, P. F., Van Bochoven, A., Visscher, P. M., & Posthuma, D. (2015). Meta-analysis of the heritability of human traits based on fifty years of twin studies. Nature genetics, 47(7), 702.

Polubriaginof, F. C. G., Vanguri, R., Quinnies, K., Belbin, G. M., Yahi, A., Salmasian, H., … Tatonetti, N. P. (2018). Disease Heritability Inferred from Familial Relationships Reported in Medical Records. Cell, 173(7), 1692-1704.e1611. doi:https://doi.org/10.1016/j.cell.2018.04.032

Tenesa, A., & Haley, C. S. (2013). The heritability of human disease: estimation, uses and abuses. Nature Reviews Genetics, 14(2), 139.

